# The differential effect of glutamine supplementation on the orthotopic and subcutaneous growth of two syngeneic murine models of glioma

**DOI:** 10.1101/2025.04.07.647607

**Authors:** Jack Maurer, Purna Mukherjee, Linh Ta, Derek C. Lee, Tomas Duraj, Juan Henao, Michael Kiebish, Devon Von Cura, Marialucia Herdocia, Thomas N. Seyfried

**Author notes:** Corresponding author, Department of Biology, Boston College, Chestnut Hill, 02467, Massachusetts, USA.

## Abstract

Glutamine serves as a major fuel source for tumor cell proliferation while simultaneously playing an essential role in maintaining gastrointestinal health and immune function. Controversy exists regarding glutamine supplementation for cancer patients undergoing chemotherapy and radiation, with concerns that it may stimulate cancer growth. The present study is the first to examine the effects of dietary glutamine supplementation (0.4g/kg/day) on the growth of malignant gliomas, which utilize large amounts of glutamine to satisfy metabolic demands. Brain bioluminescence and subcutaneous tumor volumes were used to assess the influence of glutamine supplementation on the growth of the syngeneic VM-M3 and CT-2A preclinical models of glioma. Glutamine supplementation had no significant effect on the orthotopic growth of the VM-M3 or the CT-2A gliomas when compared to non-supplemented controls. However, glutamine supplementation significantly increased tumor volume by 28% in the VM-M3 and by 166% in the CT-2A tumors when grown subcutaneously outside the central nervous system (CNS) relative to controls. Additionally, glutamine supplementation increased serum glutamine despite a localized decrease in intratumoral glutamine concentrations. Caution is warranted when considering glutamine supplementation in patients with glutamine-dependent malignancies. Further studies are needed to better understand the potential risks and benefits of glutamine supplementation in cancer therapy.

## Introduction

Glutamine is the most abundant non-essential amino acid in human physiology, with normal blood concentrations ranging from 0.4–1.0 mmol/L (1). Glutamine influences multiple biological processes including immune function, metabolism, redox balance, and homeostasis (2). Tumors exhibit an increased metabolic demand for glutamine to sustain rapid growth and proliferation *in vitro*, which suggests an interesting relationship between glutamine dependence and cancer (3,4).

Clinically, high-dose glutamine supplementation has been used alongside standard chemotherapy and radiation therapies to mitigate gastrointestinal toxicities, particularly by restoring the intestinal lining. Doses as high as 0.65 g/kg of glutamine have been administered to cancer patients with no reports of toxicity or intolerance (5). Therapeutic dosages typically range between 0.25–0.50 g/kg/day, and while select studies report amelioration of gastrointestinal side effects, they show no direct positive effects on the anti-cancer efficacy of chemotherapy or radiation (6–8). High-dose glutamine supplementation has, however, shown promise in alleviating cancer cachexia, a syndrome characterized by severe muscle and weight loss, where glutamine acts to restore muscle health and nutritional status (9,10).

More recently, preclinical studies have linked glutamine supplementation to inhibition of tumor growth through immune modulation and epigenetic mechanisms (11,12). Gabra *et al*. recently demonstrated that glutamine supplementation downregulates cancer-associated pathways in melanoma via epigenetic modifications (11). Conversely, another study showed that glutamine supplementation enhanced type-1 conventional dendritic cell (cDC1)-mediated CD8+ T cell immunity, overcoming resistance to immunotherapies and inhibiting tumor growth (12). Despite these findings, the relationship between glutamine and cancer cell proliferation in a physiological system remains complex. Glutamine is an essential amino acid for the proliferation and viability of cancer cells *in vitro* making it a critical target for cancer therapy (13). An additional study verified that human mesenchymal glioblastoma in an orthotopic mouse model exhibited high glutamine uptake and metabolic dependence when mice were infused with C^13^ labeled glutamine (14). Numerous preclinical studies have focused on depriving cancer cells of glutamine, using compounds like 6-Diazo-5-oxo-L-norleucine (DON) and JHU-083, which inhibit glutamine-dependent processes that are linked to reduced tumor growth (15,16).

The question of whether glutamine supplementation or glutamine targeting is most beneficial for cancer treatment remains unresolved. We hypothesize that the metabolic utilization of glutamine in high-grade glioma (HGG) cells may counteract the physiological benefits offered by glutamine supplementation. To test this hypothesis, we used a luciferin-luciferase bioluminescence ATP assays and tumor volume measurements to evaluate the growth of two HGG cell lines, VM-M3 and CT-2A, in both orthotopic and subcutaneous mouse models. Glutamine was supplemented through both food and water, and serum glutamine concentrations were measured to confirm intake. These cell lines, though differing in morphology and genetic backgrounds, mimic the metabolic rewiring and unrestrained proliferation characteristic of glioblastoma (GBM). This study is the first to investigate the effects of sustained dietary glutamine supplementation on the *in vivo* growth and proliferation of the VM-M3 and CT-2A brain tumors that express mesenchymal and stem cell characteristics seen in human GBM, respectively (17,18).

## Materials and Methods

### Animal Models

Mice of the VM/Dk (VM) strain were obtained as a gift from H. Fraser (University of Edinburgh, Scotland). The C57BL/6J (B6) mice were obtained originally from the Jackson Laboratory, Bar Harbor, ME. All mice used in this study were housed and bred in the Boston College Animal Care Facility using IACUC approved husbandry conditions. Male and female mice between 8–10 weeks of age were used for all studies. All animal procedures and protocols were in strict accordance with the NIH Guide for the Care and Use of Laboratory Animals and were approved by the Institutional Animal Care Committee at Boston College under assurance number A3905–01.

### Cell Lines

The VM-M3 tumor used in this study arose spontaneously in the cerebrum of an adult male mouse of the VM/Dk inbred strain. A cell line was prepared from the VM-M3 tumor. The VM-M3 tumor manifests all the invasive characteristics seen in human GBM (19). The CT-2A tumor was originally produced from implantation of 20-methylcholanthrene into the cerebral cortex of a C57BL/6J mouse and was broadly classified as a poorly differentiated highly malignant anaplastic astrocytoma and a cell line was produced from this tumor (20). Later studies showed that the CT-2A cells have characteristics of glial stem cells (18). Both cell lines are mycoplasma free. The VM-M3 and the CT-2A cell lines were transfected with a lentivirus vector (gift from Dr. Miguel Sena-Esteves) containing the firefly luciferase gene under control of the cytomegalovirus promoter (VM-M3/Fluc) to produce VM-M3/Fluc and CT-2A/Fluc cell lines. This transfection allows the cells to be tracked in the brain using bioluminescent imaging.

### Orthotopic and Subcutaneous Implantation

Tumor implantation was performed using our standard protocol (17). Mice were briefly anesthetized with 5% isoflurane in oxygen (Covetrus, Ohio, USA). The heads were disinfected with ethanol and a small incision was made in the scalp over the midline. A 3 mm^3^ burr hole was made in the skull over the right parietal region behind the coronal suture and lateral to the sagittal suture. Small (1 mm^3^) tumor fragments were implanted approximately 1.5–2.0 mm deep into the cortical region using a trocar. The skin flaps were closed with 7 mm reflex clips. The mice were placed in a warm room (25°C) until they were fully recovered. The procedure confirms 100% recovery within a few hours of implantation. The GBM tumor cells are highly invasive regardless of implantation method, and all tumor-implanted mice reach morbidity in approximately 12–15 days. The VM-M3/Fluc and CT-2A cells were implanted subcutaneously into a VM/Dk and C57BL/6J mouse as described previously (21). Tumor from donor mice was excised and chopped into tiny fragments. Each mouse received 0.2 mL of tumor fragments suspended in 0.3 mL of PBS. Mice were anesthetized with isoflurane and the tumor was implanted by subcutaneously using a 1 cc tuberculin syringe and 18-gauge needle into the right flank.

### Dietary Regiment and Body Weight

Mice were fed a standard chow diet *ad libitum* until 5 days post tumor implantation. At day 5, mice were randomly separated into control and glutamine receiving groups. L-glutamine (Cat#G3126, Sigma, MO, USA) supplementation was administered in food (25 mg glutamine/25g of powdered chow/cage/day) and water (1.0 mg/mL) until sacrifice. Each cage had ∼5 mice per cage. Total glutamine intake was estimated to be approximately 0.4g/kg per day. The average mouse weighed 25 grams and body weight was measured periodically and at the study endpoint. Glutamine was administered to orthotopic tumor bearing mice for 12-15 days and for 14-17 days in the subcutaneous models.

### Bioluminescence Imaging

The AMI HT imaging machine (Cat#A1854, Spectral Instrument Imaging, AZ, USA) was used to record the bioluminescent signal from the transfected tumor cells. For *in vivo* imaging, mice received an *i*.*p*. injection of d-luciferin (50 mg/kg) in PBS and isoflurane (5% in oxygen). Imaging times ranged from 1 to 3 min, depending on the time point. For *ex vivo* imaging, brains were removed and imaged in 0.3 mg d-luciferin in PBS. Data acquisition and analysis was performed with Living Image software (Caliper LS, MA, USA).

### Quantitative Glutamine Analysis by Ultra-High Performance Liquid Chromatography-Tandem Mass Spectrometry (UHPLC-MS/MS) and Glutamine/Glutamate-Glo Enzymatic Assay

Frozen tumor samples were homogenized with 300ul of 0.1X PBS using an Omni Bead Rupture homogenizer at 4°C. Metabolites were extracted from 100µL of tumor homogenates, or 50µL of serum, using a solid-phase extraction (SPE) 96-well plate (Isolute PLD+, Biotage, Salem, NH, USA) with 700µl of 8:1:1 acetonotrile:methanol:acetone +0.1% formic acid at -20°C. All samples included glutamine-d5 (MedChemExpress,Monmouth Junction, NJ, USA) as internal standard. Samples were dried under nitrogen and reconstituted in 100µL of 1:1 methanol:water and storage at 4°C until analysis. Absolute quantification of glutamine was performed via isotope dilution mass spectrometry. An 11-point standard curve was included in the analysis, ranging from 0.025 µg/mL up to 50 µg/mL glutamine. A weighted linear regression was applied, and concentrations were normalized using volume (serum) or total protein concentrations for each tumor, as determined using the Pierce BCA Protein Assay Kit from the original tissue homogenates. Analysis using UHPLC-MS/MS were performed on an Agilent 1290 Infinity HPLC coupled to a Thermo Q-Exactive Plus mass spectrometer as previously described (22). Serum glutamine was also quantified using the Glutamine/Glutamate-Glo™ Assay (Cat# J8021,Promega, WI, USA). Serum samples were diluted 1:10 in PBS and added to a clear 96-well plate (50 μL/well) alongside glutamine standards (0-50 μM). To measure total glutamine plus glutamate, 50 μL of Glutaminase/Inactivation Solution was added to designated wells, while parallel wells received Inactivation Solution alone for glutamate-only measurement. After 30-minute incubation at 37°C, 100 μL of Glutamate Detection Reagent was added to each well. Following 60-minute room temperature incubation, luminescence was measured. Glutamine concentration was calculated by subtracting the glutamate-only signal from the total signal using the standard curve and adjusting for the dilution factor.

### Blood Glucose Measurements

All mice were fasted for 3h before blood collection to stabilize blood glucose levels. Whole blood from the tail was placed onto the glucose strip and glucose levels (mg/dL) were measured using the Keto-Mojo monitoring system (Keto-Mojo, Napa, California).

## Results

The aim of this research was to determine if enteral glutamine supplementation could influence the growth and proliferation of two glioma cell lines *in vivo*. A secondary objective was to investigate the difference between tumors grown in the immune privileged environment of the brain versus subcutaneously in the flank. A Luciferin-luciferase based bioluminescence ATP assay or tumor volume measurements were used to assess tumor progression in the brain and flank, respectively.

### Glutamine has no significant effect on the VM-M3 or the CT-2A tumors grown orthotopically in the brain

VM-M3 and CT-2A tumor fragments were orthotopically implanted into adult VM/Dk and C57BL/6J mice respectively. Tumor take was confirmed, and glutamine supplementation began five days post-implantation. Bioluminescence was used to assess tumor progression after 12-14 days of supplementation **(Figure 1)**. Three biological replicates were performed for both cell lines to account for variability in sex, tumor implantation, and baseline bioluminescence values. Glutamine supplementation neither significantly stimulated nor inhibited orthotopic tumor growth in VM/Dk or C57BL/6J mice, though a trend toward a stimulatory effect was observed in VM-M3 **(Figure 2A-B)**. No significant sex-specific differences in response to glutamine supplementation were found **(Figure 2A-B, Experiment 2)**. Prior to animal sacrifice, serum was collected from the submandibular vein. Serum glutamine concentrations were quantified in control and glutamine supplemented mice bearing orthotopic tumors with Glutamine/Glutamate-Glo enzymatic assay (Promega, WI). Serum glutamine concentration was elevated in VM/Dk mice and significantly increased in C57BL/6J mice receiving supplementation **(Figure 2C-D)**. We did not expect glutamine concentrations to appreciably change in brain tissue due to tight regulation of amino acid concentration in CSF detailed in previous studies (23).

**Figure 1.**
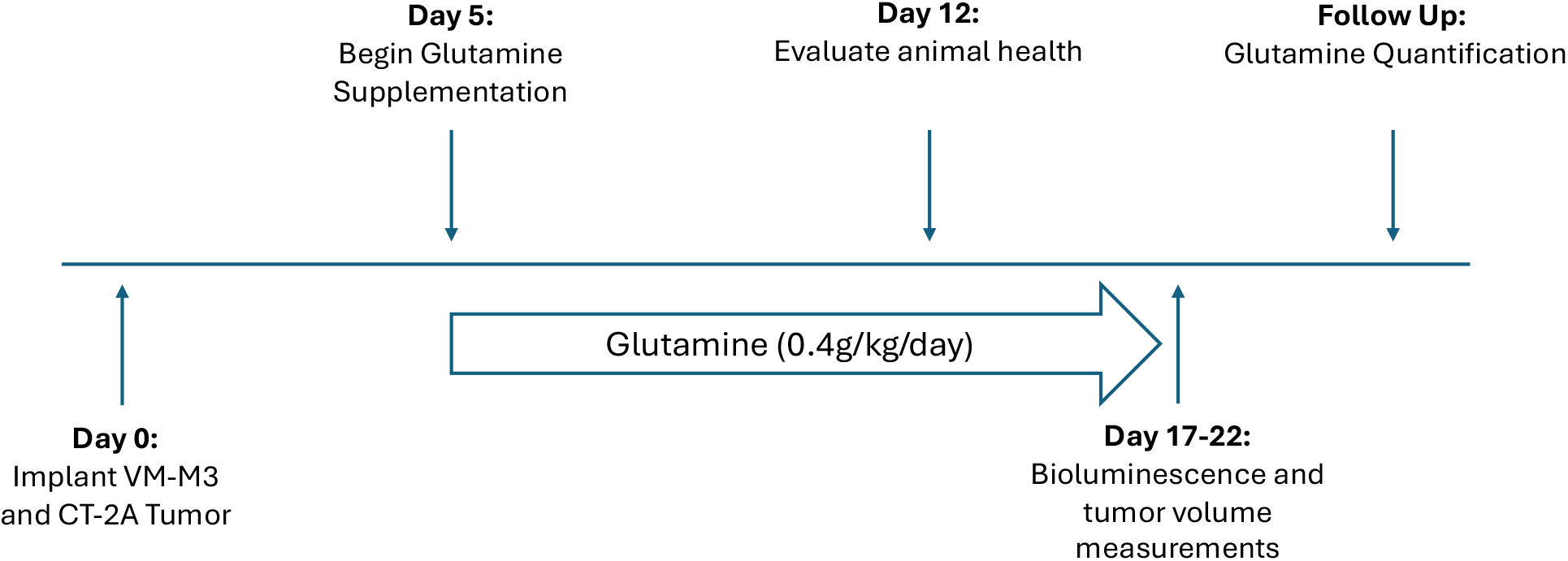
Workflow of glutamine supplementation in two syngeneic glioma models. VM/Dk and C57BL/6J mice were orthotopically implanted on day 0. On day 5, mice were divided into two dietary groups: a standard chow diet *ad libitum* (Control) or a glutamine-supplemented diet (Gln). Diet administration continued for 12–15 days, until mice became moribund. Tumor progression was monitored with *in vivo* bioluminescence imaging. In the subcutaneous model, tumor fragments were implanted into the right flank of mice and glutamine was administered for 14-17 days. Tumor volume measurements were used as the primary marker for tumor growth in subcutaneous models. Blood samples were collected from the submandibular vein, and tissue was excised and snap-frozen at -80°C for analyses. Serum and tumor tissue glutamine levels were quantified using liquid chromatography-mass spectrometry (LC/MS) and the Glutamine/Glutamate-Glo two-step enzymatic assay.

**Figure 2.**
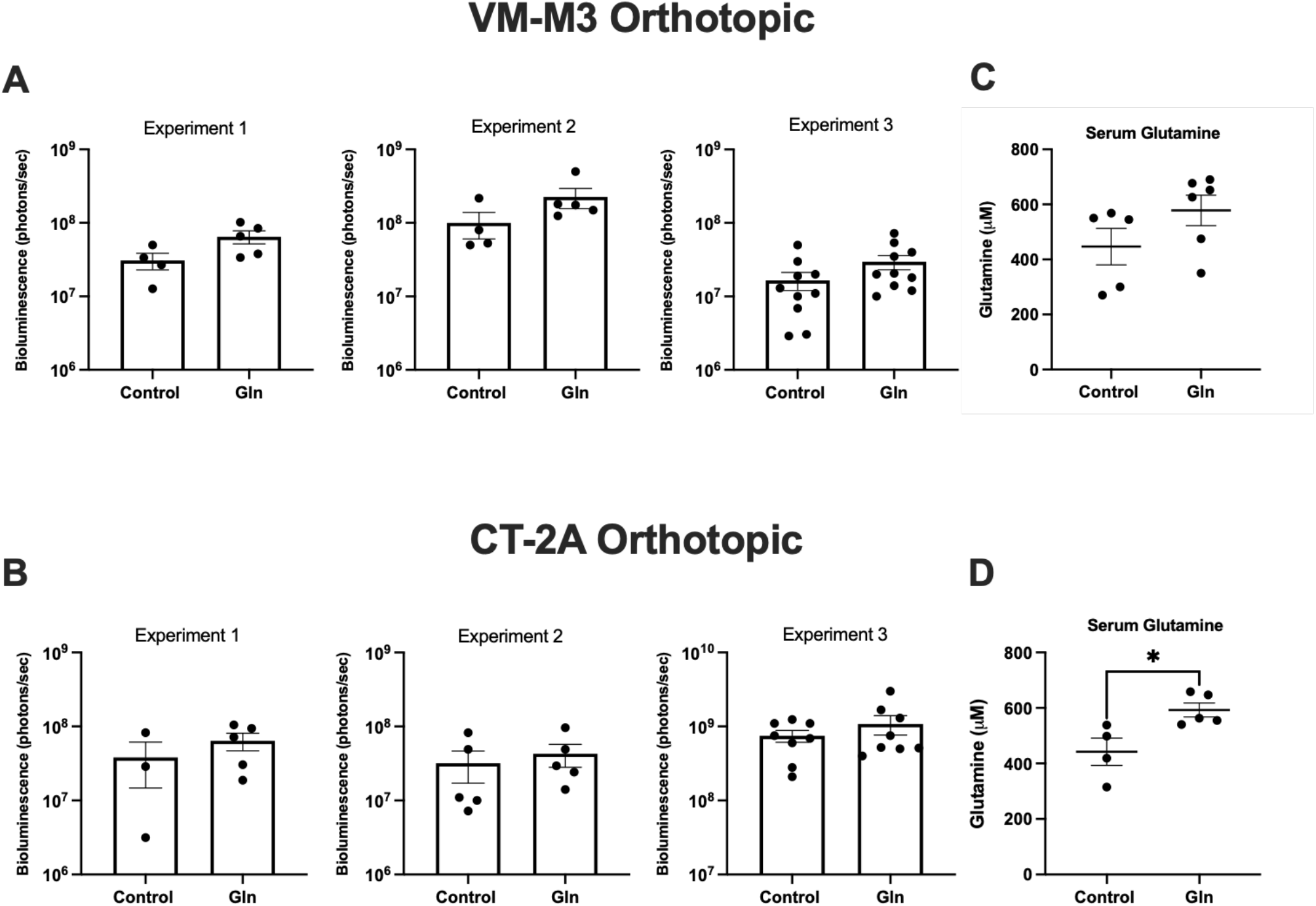
Glutamine does not affect the growth of high-grade gliomas in the syngeneic brain. **A and B**. Quantification of bioluminescence signal (photons/sec) in orthotopically implanted VM-M3 (Control n = 18, Gln n=19) and CT-2A (Control n = 16, Gln n = 18) tumors following 12-15 days of glutamine supplementation. Three biological replicates were performed with each cell line to account for variability in age, sex, and baseline bioluminescence values. Experiment 2 in both cell lines were performed with female mice. **C and D**. Serum glutamine quantification (*µ*M) in VM/Dk and C57BL/6J mice bearing orthotopic VM-M3 and CT-2A tumors respectively. All statistical comparisons were performed using the Mann-Whitney U test. Significance is indicated as follows: *p < 0.05, **p < 0.01. Data are presented as mean ± SEM.

### Glutamine stimulates growth in the VM-M3 and the CT-2A tumors when grown outside the central nervous system

To assess glutamine’s impact on tumor growth outside the brain, VM-M3 and CT-2A tumor fragments were implanted subcutaneously into VM/Dk and C57BL/6J mice and supplemented with glutamine for 14-17 days until mice became moribund. Glutamine supplementation significantly increased VM-M3 and CT-2A tumor volume **(Figure 3A-B)**. Tumor volume was chosen over bioluminescence as a marker for tumor growth for flank tumors due to its direct correlation with tumor mass and disease progression. Serum glutamine concentrations were elevated following supplementation in both models, measured with LC/MS and the Glutamine/Glutamate-Glo enzymatic assay (Promega, WI) **(Figure 3C-D)**. LC/MS analysis of intratumoral glutamine concentration in subcutaneous VM-M3 and CT-2A tumors revealed that, despite elevated serum glutamine availability, intratumoral glutamine concentrations decreased **(Figure 3E-F)**. These results suggest strong replicability between the VM-M3 and CT-2A cell lines grown orthotopically or subcutaneously in their syngeneic hosts. Preliminary blood glucose measurements indicate that glutamine supplementation has opposing effects in tumor-bearing and non-tumor-bearing C57BL/6J mice. In non-tumor-bearing mice, glutamine supplementation increased blood glucose levels over 15 days **(Supplementary Figure 1A)**. However, in CT-2A tumor-bearing mice, glutamine supplementation decreased blood glucose levels over the same period **(Supplementary Figure 1B)**. These findings suggest that tumor presence may alter systemic glucose metabolism in response to glutamine.

**Figure 3.**
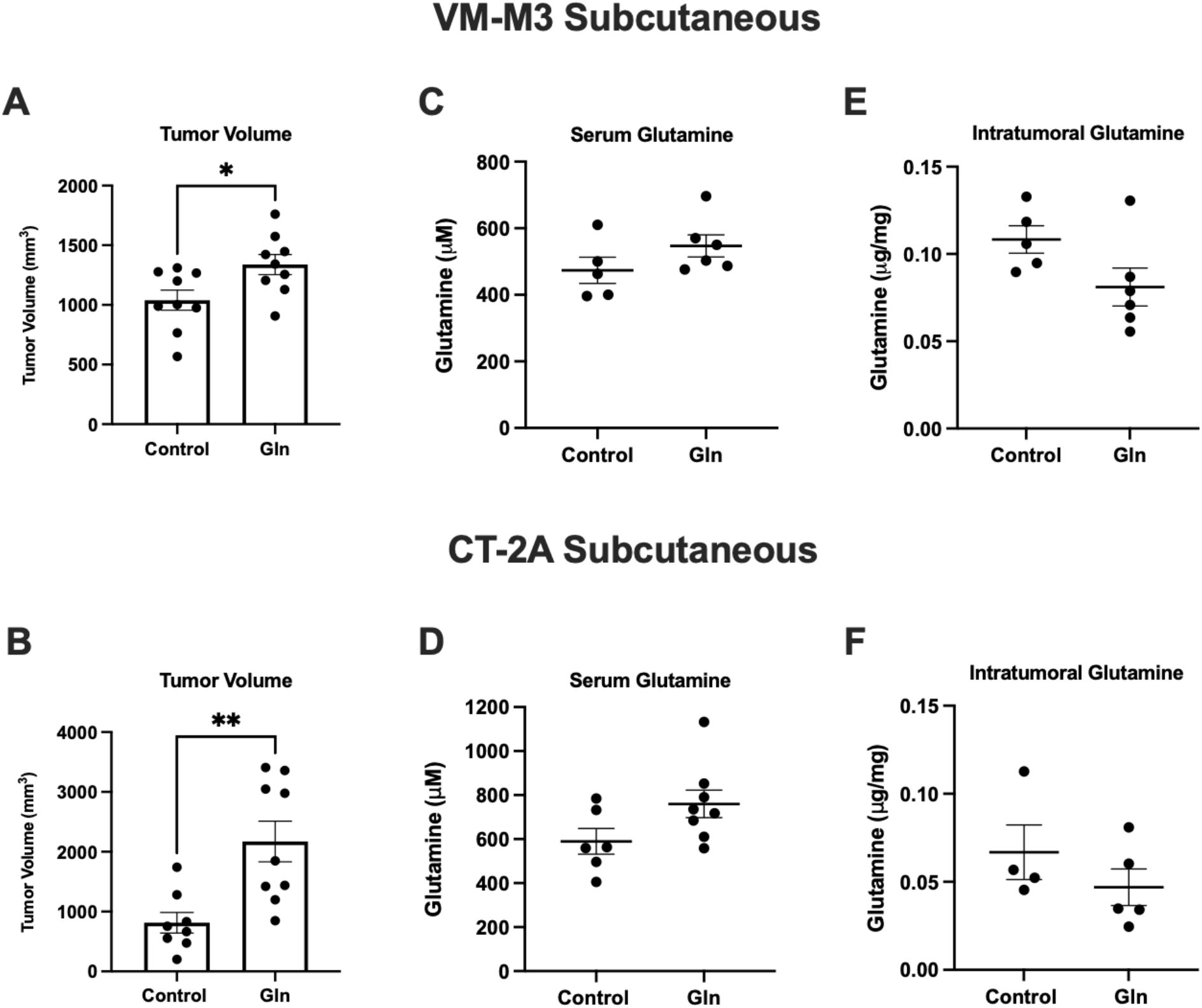
Glutamine stimulates progression of extraneural, subcutaneously grown tumors. **A and B**. Tumor volume (mm^3^) measured in subcutaneous VM-M3 (Control n=9, Gln n=9) and CT-2A (Control n=8, Gln n=9) tumors after 14-17 days of glutamine supplementation. **C and D**. Serum glutamine quantification (*µ*M) in VM/Dk and C57BL/6J mice with VM-M3 and CT-2A tumors respectively. **E and F:** Absolute intratumoral glutamine concentration (*µ*g/mg) measured with LC/MS as described in methods. Glutamine measurements were performed on n= 4-8 samples. All statistical comparisons were performed using the Mann-Whitney U test. Significance is indicated as follows: *p < 0.05, **p < 0.01. Data are presented as mean ± SEM.

## Discussion

The aim of this study was to determine if high-dose glutamine supplementation influenced the growth of two high-grade gliomas *in vivo*. Our results demonstrate that glutamine supplementation had no significant effect on orthotopic glioma growth but significantly stimulated subcutaneous tumor growth in both the VM-M3 and the CT-2A models. These findings reveal important microenvironment-dependent effects of glutamine metabolism on tumor progression that have implications for cancer patients considering glutamine supplementation. In contrast to other studies, which primarily focused on the influence of glutamine in mitigating chemotherapy- and radiation-induced gastrointestinal toxicities, our study focused on the influence of on tumor growth as a primary endpoint (6,7,24). Preclinical studies using engineered cell lines and patient-derived xenografts grown in immunocompromised mice may fail to capture the full complexity of tumor progression. In contrast, our study employed syngeneic models with intact immune systems, enabling a more complete and physiologically relevant characterization of glutamine’s effects on glioma growth within the natural tumor-host microenvironment. (25,26).

The differential response between orthotopic and subcutaneous tumors suggests that tissue microenvironment significantly influences glutamine metabolism and utilization. The lack of significant growth stimulation in brain tumors might reflect the tight regulation of glutamine levels in the glutamine glutamate cycle and by the blood-brain barrier (27,28). Previous studies have demonstrated that glutamine concentrations in the central nervous system remain carefully controlled, often equal to or lower than peripheral levels regardless of supplementation (23,27,29). This regulatory mechanism may have prevented the increased glutamine availability observed in the peripheral circulation from significantly affecting orthotopic tumor growth. Additionally, the heterogeneous glutamine metabolism across different CNS cell types may have further influenced the metabolic demands and plasticity of the brain tumors (30,31). Conversely, the significant growth stimulation observed in subcutaneous tumors suggests these systemically located malignancies directly benefit from increased glutamine availability. The decreased intratumoral glutamine concentrations despite elevated serum levels may indicate rapid glutamine consumption, supporting the glutamine addiction phenomenon described in many aggressive cancers (32). This finding aligns with glutamine’s established role as a critical metabolite for energy production, protein synthesis, and nucleotide biosynthesis in rapidly proliferating cells (13,33,34). The opposing effects of glutamine supplementation on blood glucose levels between tumor-bearing and non-tumor-bearing mice further indicate a systemic metabolic reprogramming induced by tumors. The decrease in blood glucose levels in tumor-bearing glutamine-supplemented mice suggests that tumors may increase whole-body glutamine utilization, potentially converting glutamine to glucose through gluconeogenesis or altering insulin sensitivity (35,36). These systemic effects warrant further investigation as they may impact patient metabolism during cancer therapy.

The high dose of glutamine used in our study (0.4g/kg/day) was selected based on clinical studies exploring glutamine’s therapeutic potential for managing chemotherapy-induced side effects (37,38). It is important to note that this dose substantially exceeds the minor increase in glutamine intake that might occur with dietary modifications such as high-protein diets (39). While our study demonstrates that this pharmacological dose stimulates subcutaneous tumor growth, the effects of lower, more physiological doses must be determined.

This study did not investigate the mechanism by which glutamine supplementation might increase subcutaneous tumor, as the primary goal was to document the relationship between dietary glutamine supplementation and glioma growth. However, future research should explore how glutamine might influence specific metabolic pathways, immune responses, and treatment efficacies in different tumor microenvironments. Additionally, time-course analyses of glutamine metabolism and alternative delivery methods could provide further insights into optimizing glutamine’s therapeutic potential while minimizing its tumor-promoting effects.

## Conclusion

In conclusion, our findings indicate that dietary glutamine supplementation does not inhibit high-grade glioma growth and may significantly stimulate tumor progression when grown outside the CNS. Clinicians should carefully weigh the potential benefits of glutamine supplementation against its tumor-promoting effects, particularly for patients with extraneural malignancies. Future research should focus on identifying biomarkers of glutamine dependence and developing strategies to selectively target glutamine metabolism in cancer cells while preserving its beneficial effects on normal tissues.

## Conflict of Interest

All authors state no conflict of interest.

## Author contributions

JM, PM, DCL, LT, TD, and TNS contributed to the conceptualization and design of the research. JM, PM, LT, MH performed the bulk of experiments and acquired the data. JH, MK, and DVC generated metabolite data sets. JM and TNS drafted and revised the manuscript. All authors have read and agreed to the published version of the manuscript.

## Research funding

We thank the Foundation for Metabolic Cancer Therapies, CrossFit Inc., Dr. Joseph C. Maroon, Dr. Edward Miller, The Broken Science Initiative, Children with Cancer UK, The Corkin Family Foundation, and the Boston College Research Expense Fund for their support.

## Data Availability

All data supporting the findings of this study are available from the corresponding author upon reasonable request.

## Notes

### Competing Interest Statement

The authors have declared no competing interest.

## References

1. Roth E. Nonnutritive Effects of Glutamine. J Nutr. 2008 Oct;138(10):2025S–2031S.

2. Cruzat V, Macedo Rogero M, Noel Keane K, Curi R, Newsholme P. Glutamine: Metabolism and Immune Function, Supplementation and Clinical Translation. Nutrients. 2018 Oct 23;10(11):1564.

3. Seyfried TN, Kiebish MA, Marsh J, Shelton LM, Huysentruyt LC, Mukherjee P. Metabolic management of brain cancer. Biochim Biophys Acta. 2011 Jun;1807(6):577–94.

4. Natarajan SK, Venneti S. Glutamine Metabolism in Brain Tumors. Cancers. 2019 Oct 24;11(11):1628.

5. Ward E, Picton S, Reid U, Thomas D, Gardener C, Smith M, et al. Oral glutamine in paediatric oncology patients: a dose finding study. Eur J Clin Nutr. 2003 Jan;57(1):31–6.

6. Yoshida S, Matsui M, Shirouzu Y, Fujita H, Yamana H, Shirouzu K. Effects of glutamine supplements and radiochemotherapy on systemic immune and gut barrier function in patients with advanced esophageal cancer. Ann Surg. 1998 Apr;227(4):485–91.

7. Anderson PM, Schroeder G, Skubitz KM. Oral glutamine reduces the duration and severity of stomatitis after cytotoxic cancer chemotherapy. Cancer. 1998 Oct 1;83(7):1433–9.

8. Bozzetti F, Biganzoli L, Gavazzi C, Cappuzzo F, Carnaghi C, Buzzoni R, et al. Glutamine supplementation in cancer patients receiving chemotherapy: a double-blind randomized study. Nutr Burbank Los Angel Cty Calif. 1997;13(7–8):748–51.

9. Berk L, James J, Schwartz A, Hug E, Mahadevan A, Samuels M, et al. A randomized, double-blind, placebo-controlled trial of a beta-hydroxyl beta-methyl butyrate, glutamine, and arginine mixture for the treatment of cancer cachexia (RTOG 0122). Support Care Cancer Off J Multinatl Assoc Support Care Cancer. 2008 Oct;16(10):1179–88.

10. May PE, Barber A, D’Olimpio JT, Hourihane A, Abumrad NN. Reversal of cancer-related wasting using oral supplementation with a combination of beta-hydroxy-beta-methylbutyrate, arginine, and glutamine. Am J Surg. 2002 Apr;183(4):471–9.

11. Ishak Gabra MB, Yang Y, Li H, Senapati P, Hanse EA, Lowman XH, et al. Dietary glutamine supplementation suppresses epigenetically-activated oncogenic pathways to inhibit melanoma tumour growth. Nat Commun. 2020 Jul 3;11(1):3326.

12. Guo C, You Z, Shi H, Sun Y, Du X, Palacios G, et al. SLC38A2 and glutamine signalling in cDC1s dictate anti-tumour immunity. Nature. 2023 Aug;620(7972):200–8.

13. Lee DC, Ta L, Mukherjee P, Duraj T, Domin M, Greenwood B, et al. Amino Acid and Glucose Fermentation Maintain ATP Content in Mouse and Human Malignant Glioma Cells. ASN Neuro. 2024;16(1):2422268.

14. Oizel K, Yang C, Renoult O, Gautier F, Do QN, Joalland N, et al. Glutamine uptake and utilization of human mesenchymal glioblastoma in orthotopic mouse model. Cancer Metab. 2020;8:9.

15. Seyfried TN, Flores RE, Poff AM, D’Agostino DP. Cancer as a metabolic disease: implications for novel therapeutics. Carcinogenesis. 2014 Mar;35(3):515–27.

16. Yamashita AS, da Costa Rosa M, Stumpo V, Rais R, Slusher BS, Riggins GJ. The glutamine antagonist prodrug JHU-083 slows malignant glioma growth and disrupts mTOR signaling. Neuro-Oncol Adv. 2021;3(1):vdaa149.

17. Shelton LM, Mukherjee P, Huysentruyt LC, Urits I, Rosenberg JA, Seyfried TN. A novel pre-clinical in vivo mouse model for malignant brain tumor growth and invasion. J Neurooncol. 2010 Sep;99(2):165–76.

18. Binello E, Qadeer ZA, Kothari HP, Emdad L, Germano IM. Stemness of the CT-2A Immunocompetent Mouse Brain Tumor Model: Characterization In Vitro. J Cancer. 2012;3:166–74.

19. Huysentruyt LC, Mukherjee P, Banerjee D, Shelton LM, Seyfried TN. Metastatic cancer cells with macrophage properties: evidence from a new murine tumor model. Int J Cancer. 2008 Jul 1;123(1):73–84.

20. Seyfried TN, el-Abbadi M, Roy ML. Ganglioside distribution in murine neural tumors. Mol Chem Neuropathol. 1992 Oct;17(2):147–67.

21. Shelton LM, Huysentruyt LC, Seyfried TN. Glutamine targeting inhibits systemic metastasis in the VM-M3 murine tumor model. Int J Cancer. 2010 Nov 15;127(10):2478–85.

22. Aristizabal-Henao JJ, Lemas DJ, Griffin EK, Costa KA, Camacho C, Bowden JA. Metabolomic Profiling of Biological Reference Materials using a Multiplatform High-Resolution Mass Spectrometric Approach. J Am Soc Mass Spectrom. 2021 Sep 1;32(9):2481–9.

23. Dolgodilina E, Imobersteg S, Laczko E, Welt T, Verrey F, Makrides V. Brain interstitial fluid glutamine homeostasis is controlled by blood–brain barrier SLC7A5/LAT1 amino acid transporter. J Cereb Blood Flow Metab. 2016 Nov;36(11):1929–41.

24. Muranaka H, Akinsola R, Billet S, Pandol SJ, Hendifar AE, Bhowmick NA, et al. Glutamine Supplementation as an Anticancer Strategy: A Potential Therapeutic Alternative to the Convention. Cancers. 2024 Mar 5;16(5):1057.

25. Letchuman V, Ampie L, Shah AH, Brown DA, Heiss JD, Chittiboina P. Syngeneic murine glioblastoma models: reactionary immune changes and immunotherapy intervention outcomes. Neurosurg Focus. 2022 Feb;52(2):E5.

26. Huszthy PC, Daphu I, Niclou SP, Stieber D, Nigro JM, Sakariassen PØ, et al. In vivo models of primary brain tumors: pitfalls and perspectives. Neuro-Oncol. 2012 Aug;14(8):979–93.

27. Lee WJ, Hawkins RA, Viña JR, Peterson DR. Glutamine transport by the blood-brain barrier: a possible mechanism for nitrogen removal. Am J Physiol-Cell Physiol. 1998 Apr 1;274(4):C1101–7.

28. Falkowska A, Gutowska I, Goschorska M, Nowacki P, Chlubek D, Baranowska-Bosiacka I. Energy Metabolism of the Brain, Including the Cooperation between Astrocytes and Neurons, Especially in the Context of Glycogen Metabolism. Int J Mol Sci. 2015 Oct 29;16(11):25959–81.

29. Hawkins RA. The blood-brain barrier and glutamate. Am J Clin Nutr. 2009 Sep;90(3):867S–874S.

30. Goryawala MZ, Sheriff S, Maudsley AA. Regional distributions of brain glutamate and glutamine in normal subjects. NMR Biomed. 2016 Aug;29(8):1108–16.

31. Andersen JV, Markussen KH, Jakobsen E, Schousboe A, Waagepetersen HS, Rosenberg PA, et al. Glutamate metabolism and recycling at the excitatory synapse in health and neurodegeneration. Neuropharmacology. 2021 Sep;196:108719.

32. Wise DR, Thompson CB. Glutamine addiction: a new therapeutic target in cancer. Trends Biochem Sci. 2010 Aug;35(8):427–33.

33. DeBerardinis RJ, Mancuso A, Daikhin E, Nissim I, Yudkoff M, Wehrli S, et al. Beyond aerobic glycolysis: transformed cells can engage in glutamine metabolism that exceeds the requirement for protein and nucleotide synthesis. Proc Natl Acad Sci U S A. 2007 Dec 4;104(49):19345–50.

34. Trejo-Solis C, Silva-Adaya D, Serrano-García N, Magaña-Maldonado R, Jimenez-Farfan D, Ferreira-Guerrero E, et al. Role of Glycolytic and Glutamine Metabolism Reprogramming on the Proliferation, Invasion, and Apoptosis Resistance through Modulation of Signaling Pathways in Glioblastoma. Int J Mol Sci. 2023 Dec 18;24(24):17633.

35. Stumvoll M, Perriello G, Meyer C, Gerich J. Role of glutamine in human carbohydrate metabolism in kidney and other tissues. Kidney Int. 1999 Mar;55(3):778–92.

36. Holeček M. Origin and Roles of Alanine and Glutamine in Gluconeogenesis in the Liver, Kidneys, and Small Intestine under Physiological and Pathological Conditions. Int J Mol Sci. 2024 Jun 27;25(13):7037.

37. Wang WS, Lin JK, Lin TC, Chen WS, Jiang JK, Wang HS, et al. Oral glutamine is effective for preventing oxaliplatin-induced neuropathy in colorectal cancer patients. The Oncologist. 2007 Mar;12(3):312–9.

38. Jiang H ping, Liu C an. [Protective effect of glutamine on intestinal barrier function in patients receiving chemotherapy]. Zhonghua Wei Chang Wai Ke Za Zhi Chin J Gastrointest Surg. 2006 Jan;9(1):59–61.

39. Matthews DE, Campbell RG. The effect of dietary protein intake on glutamine and glutamate nitrogen metabolism in humans. Am J Clin Nutr. 1992 May;55(5):963–70.

