## Supplementary Figures for "The differential effect of glutamine supplementation on the orthotopic and subcutaneous growth of two syngeneic murine models of glioma"

**Supplementary Figure 1.**

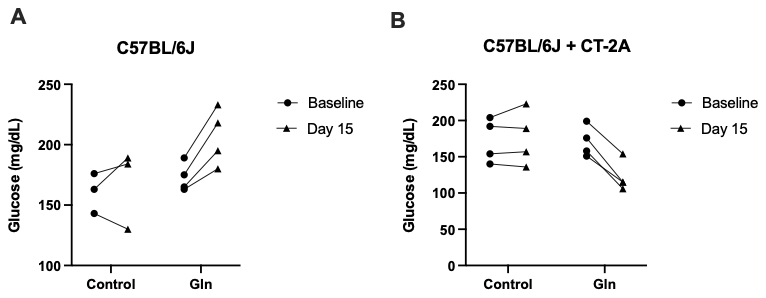

**Glutamine alters blood glucose levels in control and tumor bearing C57BL/6J mice. A.** C57BL/6J mice were fasted for three hours, and blood glucose levels (mg/dL) were measured from the tail vein using the KetoMojo glucose monitor at baseline and after 15 days of dietary intervention with or without glutamine supplementation. **B.** Mice were implanted with CT-2A tumors following baseline glucose measurements and received oral glutamine supplementation for 15 days. Blood glucose levels were monitored to assess the effects of glutamine on systemic glucose metabolism. Data are presented as absolute measurements at day 0 and day 15. 3-4 mice were used in each group.
